# High circulating elafin levels are associated with Crohn’s disease-associated intestinal strictures

**DOI:** 10.1101/739920

**Authors:** Jiani Wang, Christina Ortiz, Lindsey Fontenot, Ying Xie, Wendy Ho, S. Anjani Mattai, David Q Shih, Hon Wai Koon

## Abstract

**Objective:** Nearly 33% of Crohn’s disease (CD) patients develop intestinal strictures. Antimicrobial peptide or protein expression is associated with disease activity in inflammatory bowel disease (IBD) patients. Circulating blood cells and intestine of IBD patients have abnormal expression of elafin, a human elastase-specific protease inhibitor and antimicrobial peptide. However, the association between elafin and CD-associated intestinal stricture is unknown. We hypothesize the elafin expression in stricturing CD patients is abnormal. We determined the expression of elafin in blood, intestine, and mesenteric fat in IBD patients.

**Methods:** Human colonic and mesenteric fat tissues and serum samples were collected from the Cedars-Sinai Medical Center and UCLA, respectively.

**Results:** High serum elafin levels were associated with a significantly elevated risk of intestinal stricture in CD patients. Machine learning algorithm using serum elafin levels and clinical data identified stricturing CD patients with high accuracy. Serum elafin levels had weak positive correlation with clinical disease activity (Partial Mayo Score and Harvey Bradshaw Index) in IBD patients. Ulcerative colitis (UC) patients had high serum elafin levels, but the increase was not associated with endoscopic Mayo score. Colonic elafin mRNA and protein expression were not associated with clinical disease activity in IBD patients, while stricturing CD patients had low colonic elafin expression. Mesenteric fat in stricturing CD patients had significantly increased elafin mRNA expression, which may contribute to high circulating elafin level.

**Conclusion:** High serum elafin levels and adipose elafin expression are associated with intestinal strictures, which may help identify intestinal strictures in CD patients.

## Introduction

Intestinal stricture formation is a debilitating complication of inflammatory bowel disease (IBD) (Kugathasan, Denson et al., 2017). Around one-third of Crohn’s disease (CD) patients develop strictures (Vienna classification B2) over ten years after diagnosis (Louis, Collard et al., 2001). Currently, there is no effective approach to prevent or reverse the development of intestinal fibrosis. Anti-inflammatory agents have little to no effect on the development of intestinal fibrosis in CD patients (Kugathasan et al., 2017). Surgical resection is the last resort for severe cases. Also, diagnosis and evaluation of intestinal strictures rely on imaging and endoscopy, which are expensive and time-consuming (Bettenworth, Nowacki et al., 2016, Lenti & Di Sabatino, 2019). Thus, new diagnostic approaches to IBD-related intestinal fibrosis are needed.

Antimicrobial peptides or proteins such as serum cathelicidin, stool lactoferrin, and fecal calprotectin (FC) demonstrated clinical utilities as IBD biomarkers (Mosli, Zou et al., 2015, Tran, Wang et al., 2017). Fecal calprotectin is clinically available for assessment of IBD disease activity (Walsham & Sherwood, 2016). Cathelicidin has anti-inflammatory and anti-fibrogenic effects in colitis models (Hing, Ho et al., 2013, Ho, Pothoulakis et al., 2013, Tai, Wu et al., 2007, Yoo, Ho et al., 2015). Elafin is a small (6kDa) elastase-specific protease inhibitor with antimicrobial functions, which is primarily expressed in immune cells, intestinal tract, vagina, lungs, and skin (Shaw & Wiedow, 2011). High serum elafin levels indicate a high risk of mortality among graft vs. host disease patients (Paczesny, Braun et al., 2010). Increased serum elafin levels were also observed in other autoimmune diseases such as rheumatoid arthritis and psoriasis (Elgharib, Khashaba et al., 2019, Olewicz-Gawlik, Trzybulska et al., 2017).

A microarray study demonstrated that colonic elafin mRNA expression was increased in ulcerative colitis (UC) patients (Flach, Eriksson et al., 2006), while there is no increase of colonic elafin mRNA and protein expression in CD patients (Schmid, Fellermann et al., 2007). Zhang’s group reported the negative correlation between elafin mRNA expression in peripheral blood leukocytes and clinical disease activity (modified Mayo score and Best CD activity index score) in IBD patients (Zhang, Teng et al., 2017, Zhang, Teng et al., 2016). Although elafin is correlated with disease activity in IBD patients, the relevance of elafin in intestinal strictures is unknown. We are interested at discovering novel biomarkers for intestinal stricture because there are currently no established biomarkers for indicating intestinal strictures.

As elafin expression in circulation and intestine are altered in IBD, elafin may be one of the communication signals between different organs. A recent study suggests that mesenteric fat wrapping (creeping fat) may be associated with risk of intestinal stricture in CD patients, but the mechanism of this association has not yet been identified (Mao, Kurada et al., 2019). Elafin expression in the adipose tissue of IBD patients is unknown. We hypothesize that a link between elafin expression and intestinal fibrosis may exist. To facilitate our research, our group established a biobank by collecting and analyzing human blood, intestinal, and adipose tissue samples from patients (Sideri, Bakirtzi et al., 2015, Tran et al., 2017, Xu, Ghali et al., 2017). This study examined the expression of elafin in circulation, intestine, and mesenteric fat in non-IBD, UC, stricturing CD, and non-stricturing CD patients comprehensively. Our data suggests the utility of elafin for identifying stricturing CD patients, which is making novel progress towards better diagnostic approaches for CD-associated intestinal stricture.

## Materials and Methods

### Patients and samples

Patient-matched human colonic and mesenteric fat samples were collected from the Cedars-Sinai Medical Center (Xu et al., 2017). Human blood samples were collected from UCLA (Tran et al., 2017). All samples were collected prospectively. Serum exosomes were prepared by total exosome isolation reagent (#4478360, ThermoFisher) and quantified by bicinchoninic acid (BCA) protein assay (#23225, ThermoFisher).

Inclusion and exclusion criteria were described in our previous reports (Tran et al., 2017, Xu et al., 2017). Inclusion criteria: IBD diagnosis was confirmed by board-certified gastroenterologists. Exclusion criteria: Pregnant women, prisoners, or minors under age 18 were not included. Additionally, patients with concurrent acute infection (CMV, *C. difficile*, and tuberculosis) and malignant conditions were excluded.

The serum sample study was approved by the UCLA Institutional Review Board (protocol number IRB 12-001499 and IRB 13-001069). All samples were collected during the indicated diagnostic procedure between 2012-2015. The colonic sample study was approved by institutional review boards (Cedars-Sinai Institutional Review Board, IRBs 3358 and 23705, and UCLA Institutional Review Board IRB-11-001527). All samples were collected during the indicated diagnostic procedure between 2010-2014. Informed consent was obtained from all subjects by the Cedars-Sinai Medical Center. Separate informed consent was waived by UCLA IRB. All methods were carried out in accordance with relevant guidelines and regulations.

### ELISA

Human colonic tissues were homogenized in RIPA buffer with a protease inhibitors cocktail (sc-24948, Santa Cruz Biotechnology). Human sera were diluted ten-fold with reagent diluent and added to the ELISA plates. Measurement of elafin was performed using an ELISA kit (DY1747 R&D Systems) as described previously (Koon, Shih et al., 2011). Measurement of serum cytokines was performed using multiplex ELISA (human 27-plex #m500kcaf0y, Bio-Rad).

### Cell Cultures

Human CCD-18Co intestinal fibroblasts (ATCC) (2 × 10^6^ cells/plate) were cultured in minimal essential medium Eagle’s medium (MEM) containing 10% fetal bovine serum and 1% penicillin-streptomycin (Invitrogen) (Xu et al., 2017, Yoo et al., 2015). Serum-starved CCD-18Co cells were treated with 15μg/ml of anti-elafin neutralizing antibody (AF1747, R&D Systems) or control antibody (AB-108-C, R&D Systems), followed by exposure to human sera from normal, UC, stricturing CD, and non-stricturing CD patients (100μl/ml).

CCD18Co fibroblasts in MEM and primary human peripheral blood mononuclear cells (PBMCs) in mononuclear cell medium (C-28030, Promocell) were incubated with 100μg/ml of human serum exosomes for 24 hours. Human serum exosomes were obtained from 12 patients per group. PBMCs were obtained from a healthy donor (C-12907, Promocell). At the end of the experiments, the treated PBMCs were centrifuged, and the cell pellets were used for RNA extraction. The treated fibroblasts were collected for RNA extraction.

### Histological evaluation of intestinal injury

We prepared paraffin-embedded sections of each human colonic biopsies at tissue processing core laboratory (TPCL) at UCLA. Paraffin-embedded sections were cut at 4μm thickness, and H&E staining of the tissue sections was performed as described previously (Xu et al., 2017, Yoo et al., 2015). Microphotographs were recorded at multiple locations, and blindly scored by two investigators (Xu et al., 2017, Yoo et al., 2015). Histology score of human colonic tissues was evaluated using a previously reported approach (D’Haens, Geboes et al., 1998).

### Elafin immunohistochemistry

Elafin immunohistochemistry of human colonic and mesenteric fat tissues was performed by TPCL. Paraffin was removed with xylene. The sections were then rehydrated through graded ethanol. Endogenous peroxidase activity was blocked with 3% hydrogen peroxide in methanol for 10 minutes. Heat-induced antigen retrieval (HIER) was carried out for all sections in 0.01M citrate buffer, pH = 6 using a Biocare decloaker at 95°C for 25 minutes. After treatment with blocking buffer (2% BSA) for 1 hour, the slides were then incubated overnight at 4°C with rabbit polyclonal to elafin in 2% BSA at 1:100 dilution (Sigma, HPA017737). The signal was detected using the rabbit horseradish peroxidase EnVision kit (DAKOCytomation, K4003). This secondary antibody kit was directly applied to the slides without dilution. All sections were visualized with the diaminobenzidine reaction and counterstained with hematoxylin. Images were taken with a Zeiss AX10 microscope in a blind manner.

### Quantitative real-time RT-PCR

Total RNA was isolated by an RNeasy kit (#74104, Qiagen) and reverse transcribed into cDNA by a high-capacity cDNA RT kit (#4368813, ThermoFisher). Quantitative PCR reactions were run with Fast Universal PCR master mix (#4352042, ThermoFisher) in a Bio-Rad CFX384 system (Koon et al., 2011). The mRNA expression was determined by using cataloged primers (ThermoFisher) for human collagen COL1A2 (Hs01028956_m1), alpha smooth muscle actin ACTA2 (Hs00426835_g1), and elafin PI3 (Hs00160066_m1). Relative mRNA quantification was performed by comparing test groups and normal control group, after normalization with endogenous control gene human 18S (Hs99999901_s1).

Preliminary screening for the presence of serum exosomal miRNAs was determined using a miScript human miFinder PCR array (MIHS-001Z, Qiagen). RNA was converted to cDNA using miScript RT kit (218060). PCR reactions were performed with miScript SYBR Green PCR kit (218073). Since many miRNAs in the PCR arrays were undetectable in serum exosomes, we selected the detectable miRNAs and determined their relative expression using miRCURY LNA miRNA PCR assays. RNA was converted to cDNA using miRCURY LNA RT kit (339340) and PCR reactions were run with miRCURY LNA SYBR Green PCR kit (339346). The miRNA expression was detected using Qiagen miRCURY PCR assays (miR29c-3p YP00204729 and miR205-5p YP00204487). Relative miRNA quantification was performed by comparing test groups and normal control group, after normalization with housekeeping miRNA (RNU1A1). Measurement of miR29c-3p and miR205-5p in PBMCs, mesenteric fat, and colonic tissues was determined by miRCURY LNA PCR assays. All miRNA-related reagents were purchased from Qiagen.

The fold changes are expressed as 2^∆∆Ct^. Fold-change values greater than one indicates a positive- or an up-regulation, and the fold-regulation is equal to the fold-change. Fold-change values less than one indicate a negative or down-regulation, and the fold-regulation is the negative inverse of the fold-change.

### Power analysis

Colonic sample study: At least 30 patients per group were required to achieve a statistically significant difference of colonic elafin mRNA expression between control (1.39 fold), UC (12 fold), and CD (4 fold) patients with standard deviation = 2.55, alpha = 0.5, and power = 0.8. The cohort consisting of 40 non-IBD, 52 UC, and 43 CD patients satisfied this requirement.

Serum sample study: At least 30 patients per group were required to achieve a statistically significant difference of serum elafin levels between control (7939pg/ml), UC (12987pg/ml), and CD (12344pg/ml) patients with standard deviation = 4860, alpha = 0.5, and power = 0.8. The combined dataset from the two serum cohorts yielded 70 control, 80 UC, and 95 CD patients total that satisfied this requirement.

### Statistical analysis

Colonic elafin mRNA and protein expression were arranged in low-to-high order. The entire range of data was divided into three equal tertiles (⅓, ⅓, ⅓). Serum elafin concentrations were arranged in order from low to high. We compared the performance of multiple cut-off points of elafin levels at each disease parameter for optimization of test performance. After many calculations using various cutoff points, the optimized universal cut-off points yielding the highest area under the curve (AUC) values in receiver operating characteristic (ROC) curves (most accurate) were shown in this study. Calculation of prevalence of the disease, sensitivity, specificity, positive predictive value, negative predictive value, and relative risk was described previously (Tran et al., 2017). AUCs of ROC curves were calculated online (easyROC web-tool, www.jrocfir.org, and Microsoft Azure Machine Learning Studio). Unpaired Student’s *t*-tests were used for two-group comparisons of continuous data (GraphPad QuickCalcs) online. One-way ANOVAs with Tukey Honestly Significant Difference *post-hoc* tests were used for multiple-group comparisons (Statpages) online. Bar graphs and scatter plots were made using Microsoft Excel. Equations and R^2^ values in scatter plots were generated by Microsoft Excel. Results were expressed as mean +/− SEM. Significant *p* values are shown in each figure.

### Machine learning algorithm for indicating the presence of stricturing in CD patients

The combined CD cohort dataset containing 67 CD patients in CSV file format was loaded into the Microsoft Azure Machine Learning Studio. The dataset included serum elafin level and 14 clinical parameters, i.e., patient’s age at blood collection (number), years of disease duration (number), CRP (number), ESR (number), HBI (number), count of IBD-related surgery (number), gender (male or female), smoking habit (yes or no), use of biologics (yes or no), use of steroid (yes or no), use of immunomodulator (yes or no), use of aminosalicylate (yes or no), presence of fistula (yes or no), and presence of stricture (yes or no). The choice of clinical data inclusion was based on repeated optimization. The machine learning algorithm regarded these data are relevant for the accurate indication of strictures. The entire dataset was split into 50% for training and 50% for evaluation. The trained model was built on a two-class decision forest algorithm. The algorithm utilized default parameters including bagging resampling method, single parameter create trainer mode, 8 decision trees, 32 maximum depth of the decision trees, 128 random splits per node, and 1 minimum number of samples per leaf node. The scored dataset showed score probability (0-0.5 indicates no stricture, 0.51-1.0 indicates stricture), scored labels (yes or no stricture), and AUC values of ROCs.

## Results

### High serum elafin levels indicated an elevated risk of stricture in CD patients

Baseline characteristics, disease locations, and medication use of the two serum sample cohorts are shown in Supplementary Table 1. The combined dataset had been used in our previous cathelicidin-IBD biomarker study (Tran et al., 2017). The detected serum elafin levels in nanogram per milliliter range were similar to the findings of other groups (Olewicz-Gawlik et al., 2017, Paczesny et al., 2010). UC patients had significantly higher serum elafin levels than control patients (Figure 1A). There was a trend of increased serum elafin levels in CD patients, but the difference was not statistically significant (Figure 1A). Serum elafin levels were directly proportional to the Harvey Bradshaw Index (HBI) in CD patients (Figure 1B), but linear regression analysis suggests the correlation was weak (low R^2^ value).

**Figure 1.**
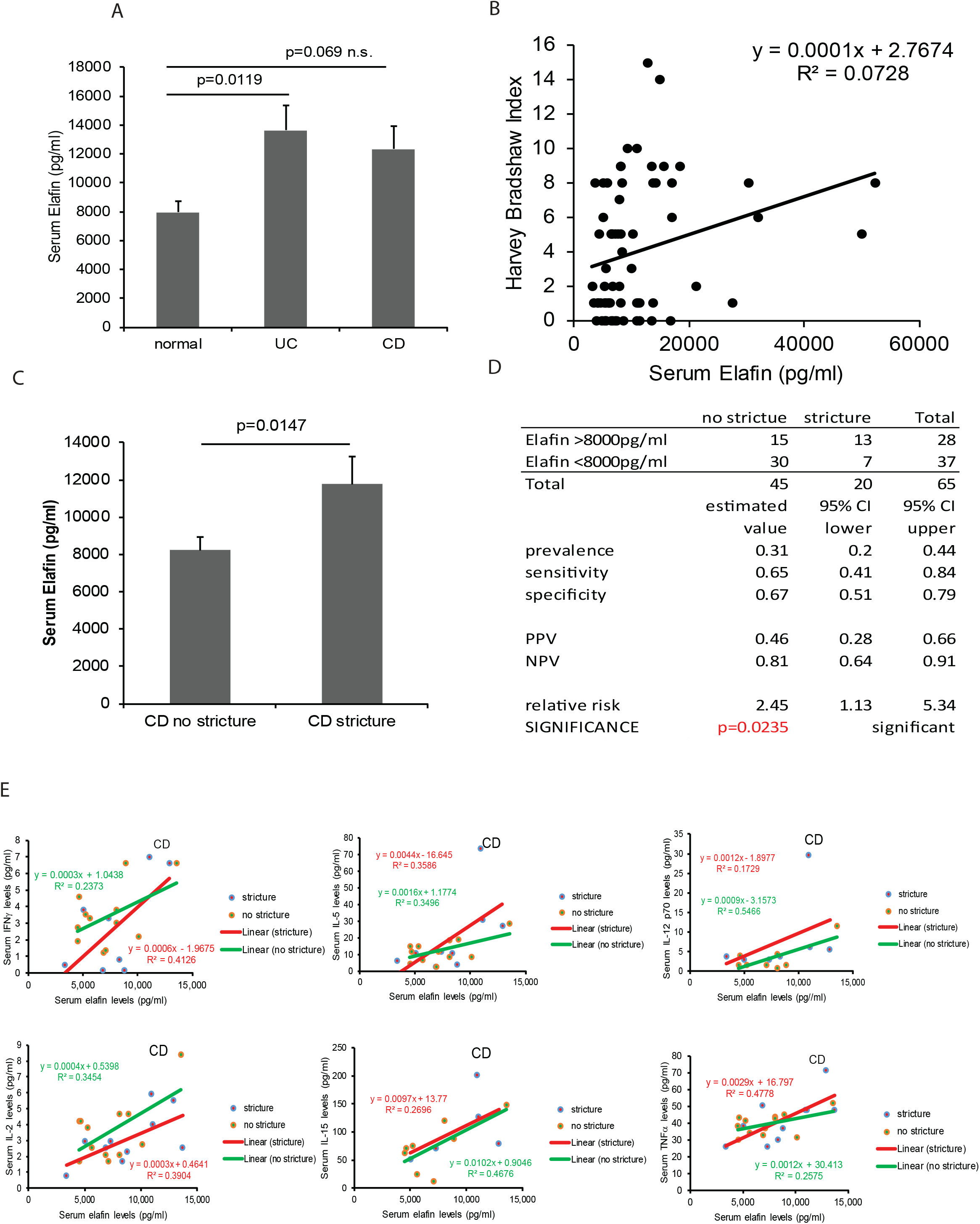
Circulating elafin levels are increased in IBD patients. (A) Serum elafin levels of normal, UC, and CD patients. Multiple group comparisons were done by one-way ANOVA. (B) Scatter plot shows the moderate correlation between serum elafin levels and clinical disease activity (Harvey Bradshaw Index) in CD patients. (C) The stricturing CD patients had significantly higher serum elafin levels than non-stricturing CD patients. Two-group comparison was done by Student’s t-test. (D) Prevalence, sensitivity, specificity, positive predictive value, negative predictive value, and relative risk of elafin test for indicating intestinal stricture in CD patients. (E) Scatter plot shows the positive correlation between serum elafin levels and serum cytokine levels in CD patients with or without intestinal strictures. Only the elafin-dependent cytokines were shown.

Stricturing CD patients had significantly higher serum elafin levels than non-stricturing CD patients (Figure 1C). The high elafin group had a significantly higher relative risk (RR = 2.45) than the low elafin group in developing intestinal strictures (Figure 1D). However, the serum elafin levels in CD patients with and without fistulas were similar, suggesting that serum elafin levels are not associated with the occurrence of fistulas in CD patients (Supplementary Figure 1A).

Multiplex ELISA of 27 cytokines indicated that serum elafin levels were positively correlated with serum IFNγ, IL-5, IL-12 p70, IL-2, IL-15, and TNFα levels in CD patients with and without the presence of intestinal strictures (Figure 1E). Twenty-one other detectable serum cytokines did not correlate with serum elafin levels (data not shown). There is no association between elafin levels, age (A1-3), disease location (L1-4), and use of medication at the time of blood collection (Supplementary Table 2).

### Machine learning algorithm improves the accuracy of elafin for indicating strictures in CD patients

To evaluate whether circulating elafin alone is a good indicator for intestinal stricture, we determined its accuracy with receiver operating characteristic (ROC) analysis. Serum elafin alone is moderately accurate for indicating stricture in CD patients (AUC=0.657 using elafin alone) (Figure 2A). We utilized machine learning to develop an algorithm for indicating the presence of intestinal strictures in CD patients (Figure 2B). The optimized trained model using serum elafin levels and commonly available clinical data together is much more accurate than those using either elafin or clinical data alone for indicating the presence of stricture (AUC = 0.917 using combined data; 0.742 using clinical data alone) (Figure 2C). Therefore, a combination of high serum elafin level and other characteristics are strongly associated with the presence of stricture in CD patients.

**Figure 2.**
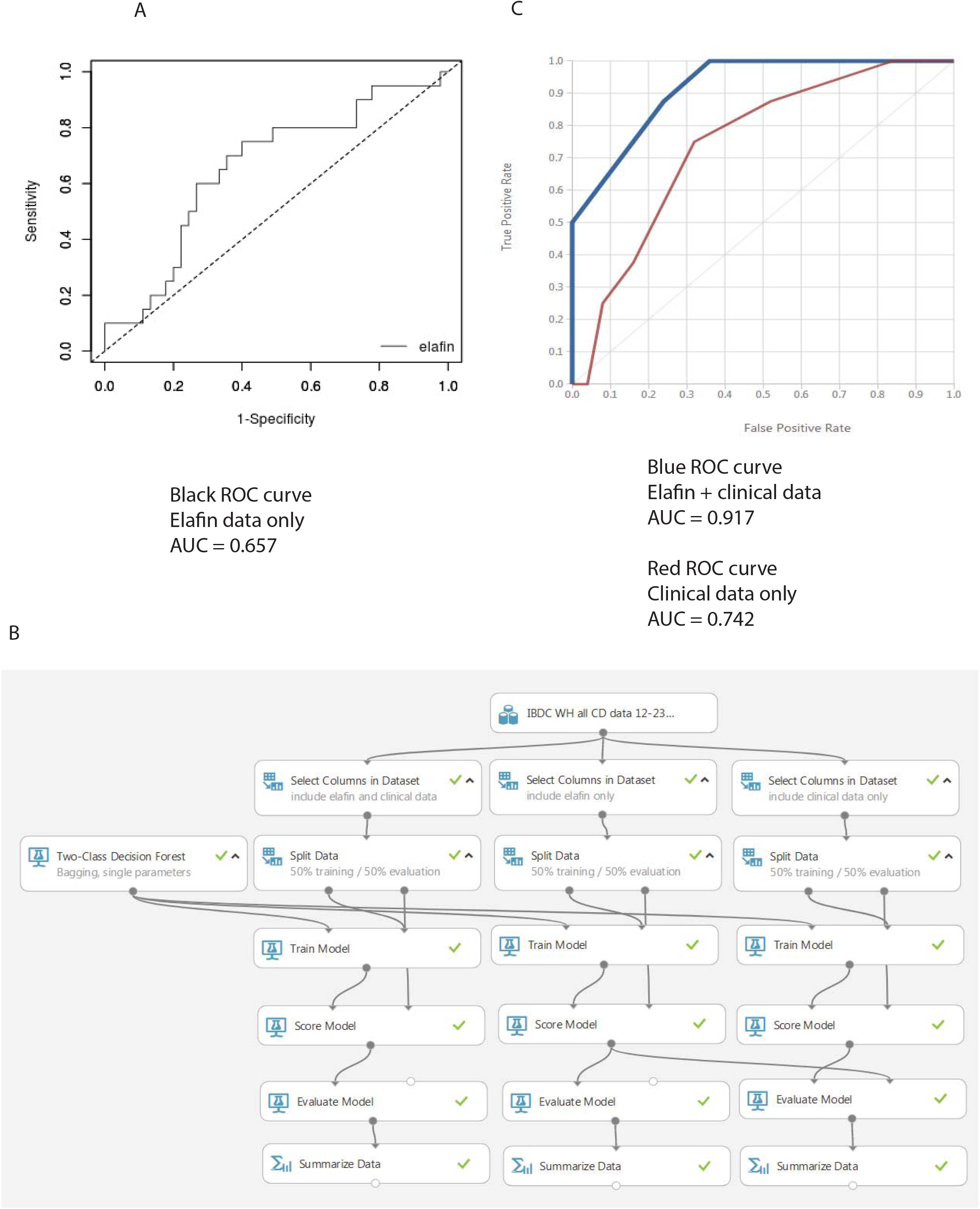
A combination of serum elafin levels and clinical data indicates the presence of stricture accurately. (A) ROC curves and AUC values show the accuracy of using elafin data alone for intestinal stricture identification among CD patients. The analysis was performed by easyROC web-tool. Cutoff elafin level is 8000pg/ml. (B) The flowchart of Microsoft Azure machine learning algorithms for indicating intestinal strictures in CD patients. (C) ROC curves and AUC values show the accuracy of using clinical data with or without elafin data for intestinal stricture identification among CD patients.

### Serum elafin levels are not correlated with endoscopic disease activity in UC patients

On the other hand, high serum elafin levels had poor positive correlation with increased Partial Mayo Score (PMS) in UC patients, as indicated by low R^2^ value (Figure 3A). Serum elafin levels had no association with Mayo Endoscopic Score in the same set of UC patients (Figure 3B). There is no association between elafin levels, disease location (E1-3), and use of medication at the time of blood collection (Supplementary Table 3). Interestingly, serum elafin levels were positively correlated with IFNγ and IL-5 levels in UC patients (Figure 3C-D). Twenty-five other detectable serum cytokines did not correlate with serum elafin levels (data not shown).

**Figure 3.**
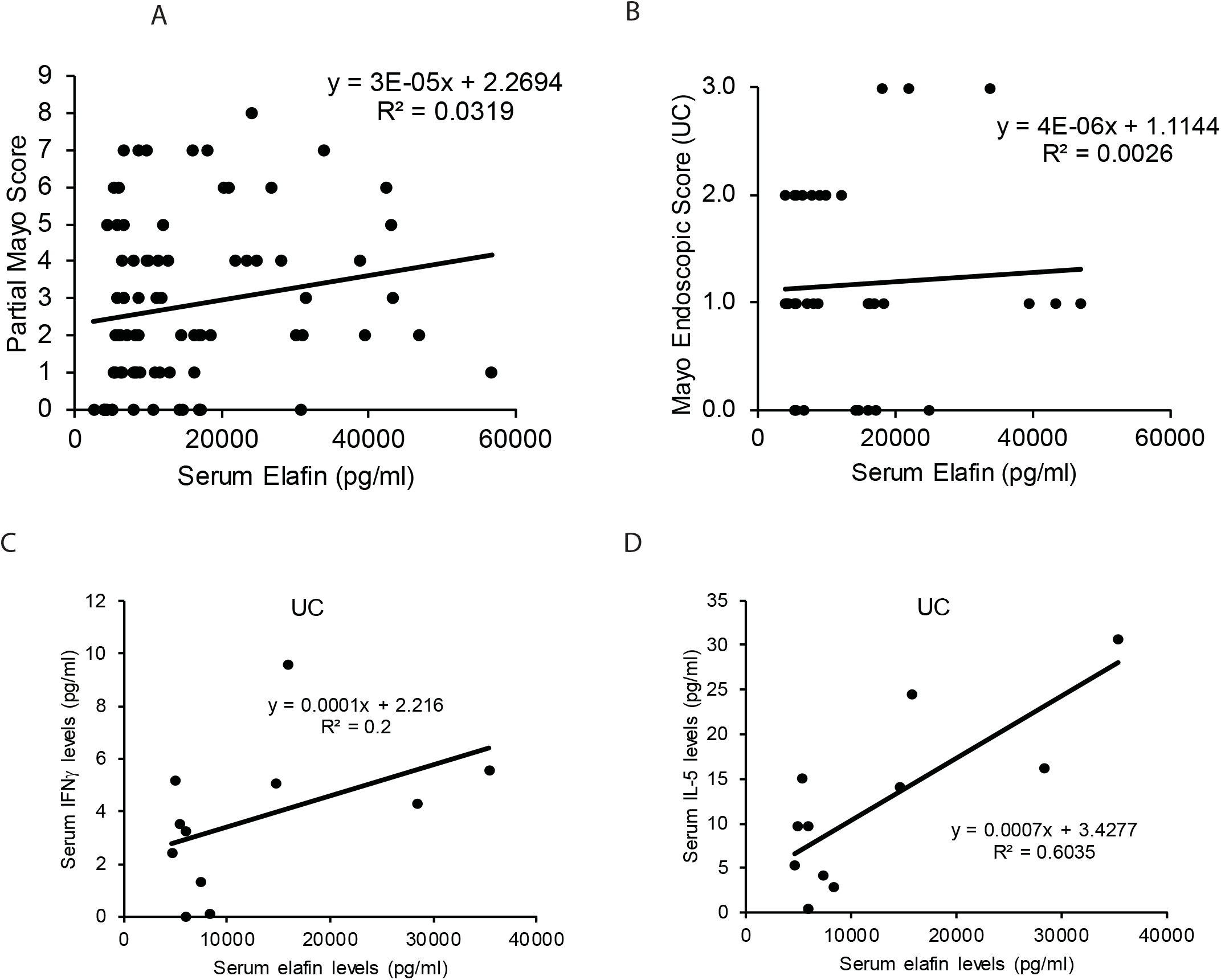
High circulating elafin levels are moderately correlated with disease activity in UC patients. (A) Scatter plot shows the positive correlation between clinical disease activity (Partial Mayo Score) and serum elafin levels in UC patients. (B) Scatter plot shows no correlation between serum elafin levels and endoscopic disease activity in UC patients. (C-D) Scatter plots show the positive correlation between serum elafin levels and serum cytokine levels in UC patients.

### Colonic elafin mRNA and protein expression were low in stricturing CD patients

Baseline characteristics of the colonic tissue cohort are shown in Supplementary Table 3 (Xu et al., 2017). Consistent with a previous study (Flach et al., 2006), UC patients had significantly higher colonic elafin mRNA and protein expression than control non-IBD patients (Figure 4A-B). Increased colonic elafin mRNA expression was positively correlated with increased colonic elafin protein expression in both UC and CD patients (Figure 4C). CD patients with stricture had significantly lower colonic elafin mRNA and protein expression than those without stricture (Figure 4D-E).

**Figure 4.**
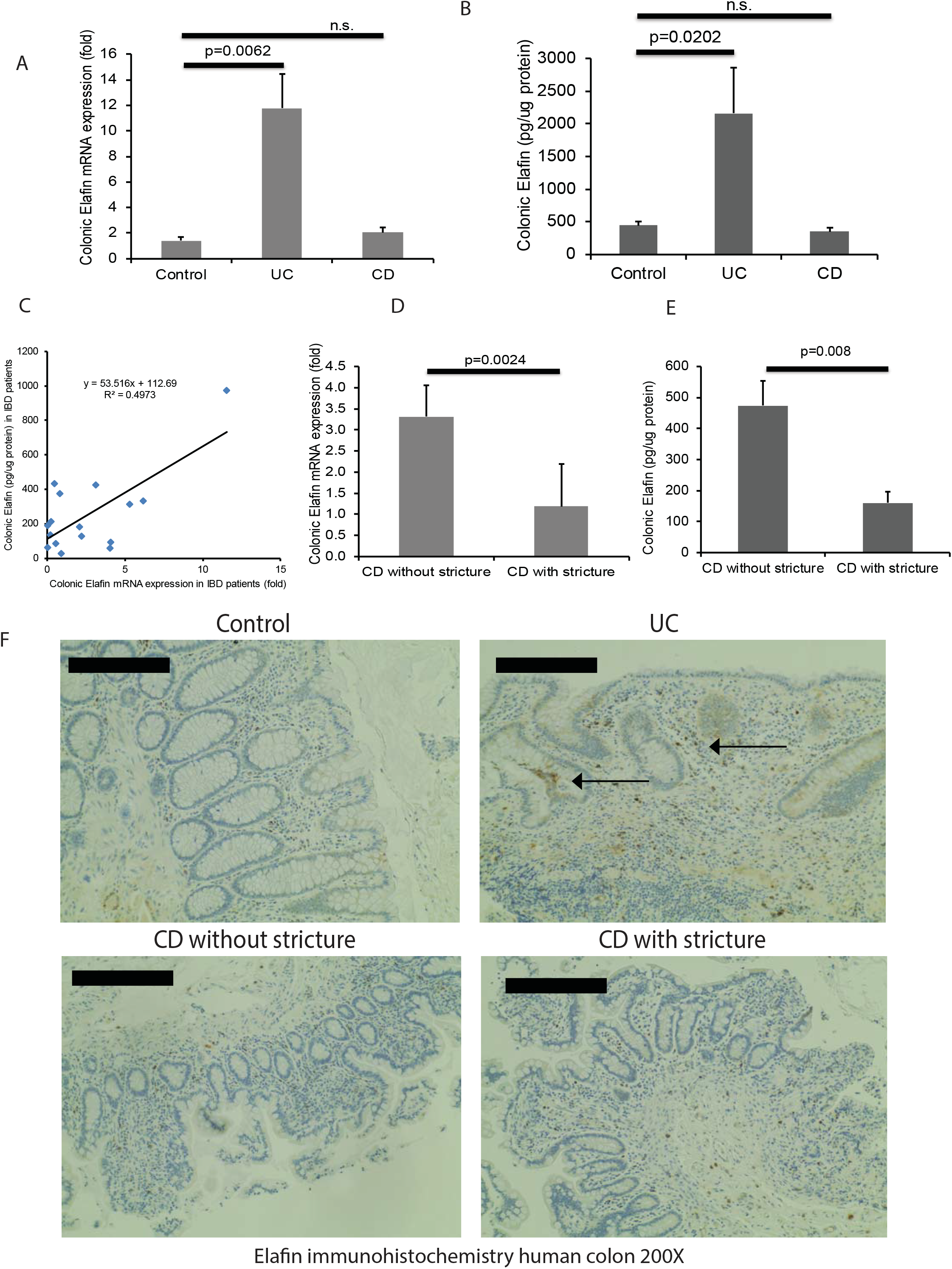
Colonic elafin mRNA and protein expression are reduced in stricturing CD patients. (A) Colonic elafin mRNA expression in non-IBD control, UC, and CD patients. (B) Colonic elafin protein expression in IBD patients. Multiple group comparison between control, UC, and CD patients were done by one-way ANOVA. (C) Scatter plots show the positive correlation of elafin mRNA and protein expression in IBD patients. (D-E) Colonic elafin mRNA and protein expression in stricturing CD and non-stricturing CD patients. Two-group comparison between CD with stricture and CD without stricture was done by Student’s t-test. (F) Immunohistochemistry of elafin in human colonic tissues. Arrows show the elafin protein in mucosal epithelial layers and lamina propria in UC patients.

Elafin protein expression in the colonic tissues of control patients was weak (Figure 4F). Elafin immunoreactivity was found in the colonic mucosa and lamina propria of UC patients (Figure 4F). Consistent with another study (Schmid et al., 2007), colonic elafin protein expression was low in CD patients with and without stricture (Figure 4F).

Collagen (COL1A2) and vimentin (Vim) are major extracellular matrix and fibroblast markers in stricture, respectively. We found that colonic elafin mRNA and protein expression were inversely proportional to colonic collagen and vimentin mRNA expression in CD patients (Figure 5A-D). Therefore, CD patients with low elafin expression tend to have increased colonic expression of collagen and accumulation of fibroblasts.

**Figure 5.**
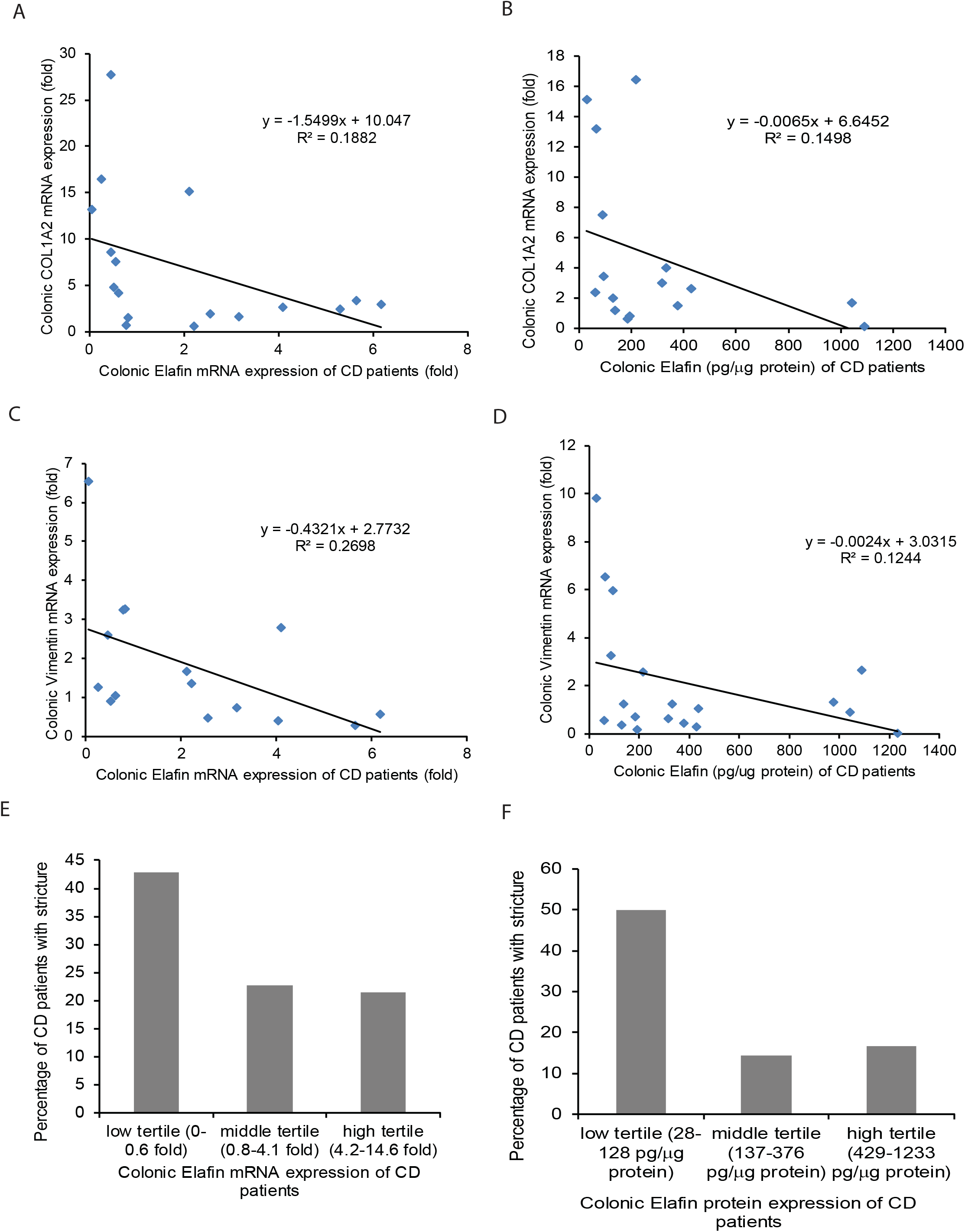
Colonic elafin mRNA and protein expression are negatively correlated with colonic fibrogenic gene mRNA expression in CD patients. (A-B) Scatter plot shows the negative correlation between colonic elafin mRNA and protein expression and colonic collagen COL1A2 mRNA expression in CD patients. (C-D) Scatter plot shows the negative correlation between colonic elafin mRNA and protein expression and colonic vimentin Vim mRNA expression in CD patients. (E-F) Percentage of intestinal stricture in CD patients assorted by colonic elafin mRNA and protein expression. Low elafin expression groups had a higher percentage of strictures than high elafin expression groups.

When the entire CD patient cohort was divided into tertiles of colonic elafin mRNA and protein expression, the low tertile tended to have a higher incidence of intestinal stricture than middle and high tertiles (Figure 5E-F). This evidence suggests that CD patients with low colonic elafin expression are prone to intestinal strictures.

Colonic elafin mRNA and protein expression were not associated with clinical disease activity in UC and CD patients (Supplementary Figure 3A-D). Colonic elafin mRNA expression had a modest negative correlation with histology score of the colonic tissues in UC and CD patients (Supplementary Figure 3E-F). The colonic elafin mRNA and protein expression had no association with age at biopsy collection, gender, use of medication, or duration of diseases in both UC and CD patients (data not shown).

### Mesenteric fat is a source of circulating elafin in stricturing CD patients

Since stricturing CD patients had high serum elafin levels and low colonic elafin expression, we continued to discover the source of elafin. Stricturing CD patients have a higher visceral to subcutaneous fat area ratio than non-IBD patients (Erhayiem, Dhingsa et al., 2011). Stricturing CD patients also have a higher visceral fat/total fat mass ratio than non-stricturing CD patients (Buning, von Kraft et al., 2015). Mesenteric fat may be a potential source of elafin.

Sticturing CD patients had significantly higher mesenteric fat elafin mRNA expression than control and non-stricturing CD patients (Figure 6A). Interestingly, our patient-matched biopsy collection indicates that mesenteric fat elafin mRNA expression is positively correlated with the colonic fibrogenic factor mRNA expression and negatively correlated with the colonic elafin protein expression in CD patients (Figure 6B-C). Immunohistochemistry indicated that elafin protein expression in mesenteric fat of stricturing CD patients was much higher than those in non-IBD, UC, and non-stricturing CD patients (Figure 6D). At high magnification (400X), the elafin-positive signal is located around adipocytes (Supplementary Figure 4A).

**Figure 6.**
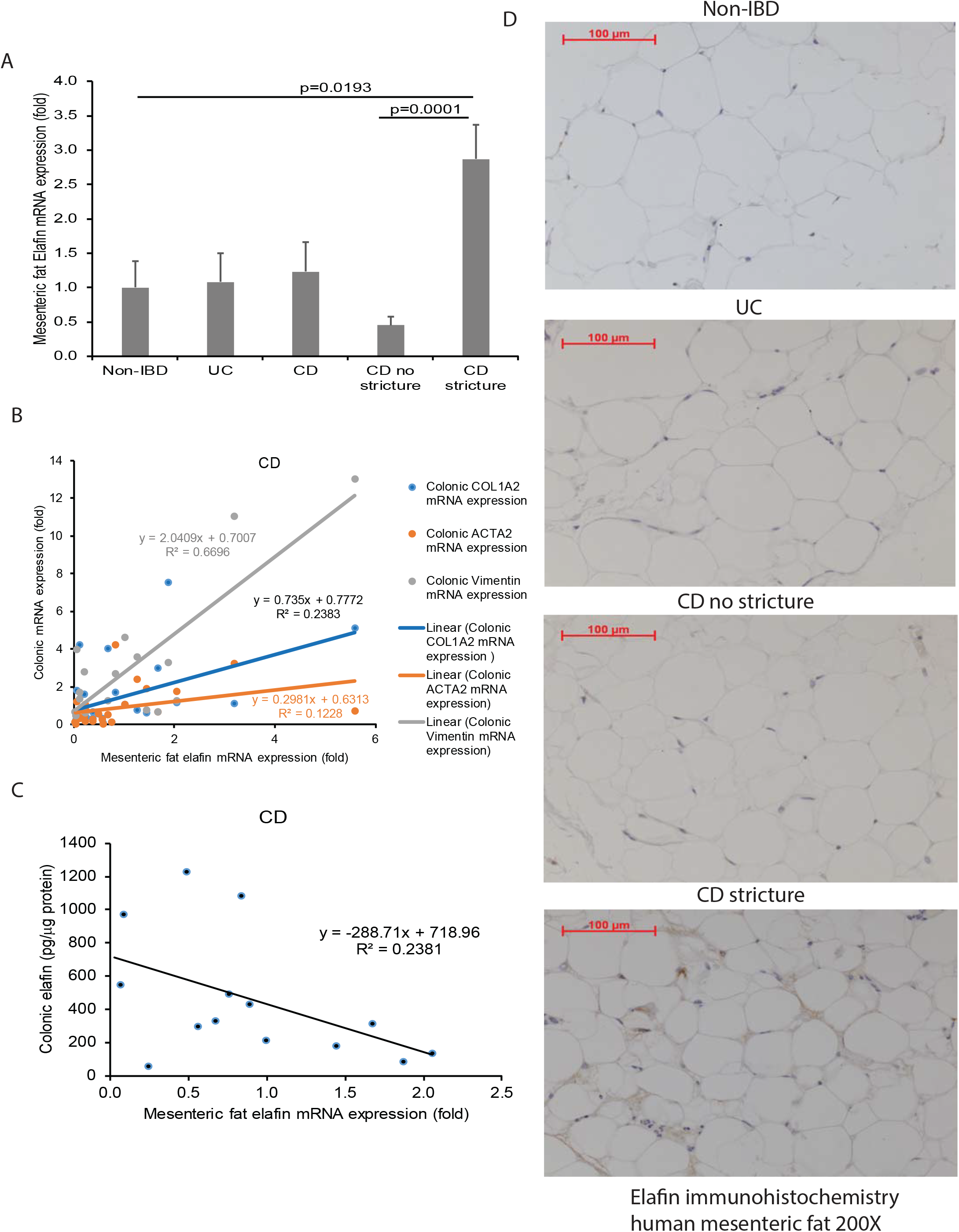
Mesenteric fat in stricturing CD patients expresses elafin. (A) Mesenteric fat elafin mRNA expression in non-IBD, UC, and CD patients. Multiple comparisons between control, UC, and CD patients were done by one-way ANOVA. Two-group comparison between CD with stricture and CD without stricture was done by Student’s t-test. (B) Scatter plot shows the positive correlation between mesenteric fat elafin mRNA expression and colonic fibrogenic gene mRNA expression in CD patients. (C) Scatter plot shows the negative correlation between mesenteric fat elafin mRNA expression and colonic elafin protein expression in CD patients. (D) Immunohistochemistry of elafin in human mesenteric fat tissues at 200X magnification. Elafin (as shown by brown color) protein expression was strong in mesenteric fat in stricturing CD patients. Four patients per group.

### Circulating exosomes, but not elafin affect fibrogenesis directly

To determine whether the circulating elafin regulates fibrogenesis, we treated the human intestinal CCD-18Co fibroblasts with human sera and their derived exosomes (Figure 7A-B). Sera from stricturing CD patients, but neither from healthy control nor non-stricturing CD patients, significantly increased collagen mRNA expression in the CCD-18Co cells (Figure 7A). Neutralization of elafin with anti-elafin antibody also did not alter the collagen mRNA expression in fibroblasts exposed to serum samples from stricturing CD patients (Figure 7A). On the other hand, serum exosomes from stricturing CD patients, but neither from control nor non-stricturing CD patients, significantly increased collagen and alpha-smooth muscle actin (ACTA2) mRNA expression in the CCD-18Co cells (Figure 7B). Therefore, circulating elafin may be a surrogate biomarker of stricture because endogenous elafin does not affect fibrogenesis directly. The serum exosomes from stricturing CD patients contain pro-fibrogenic mediators that induce collagen expression in fibroblasts.

**Figure 7.**
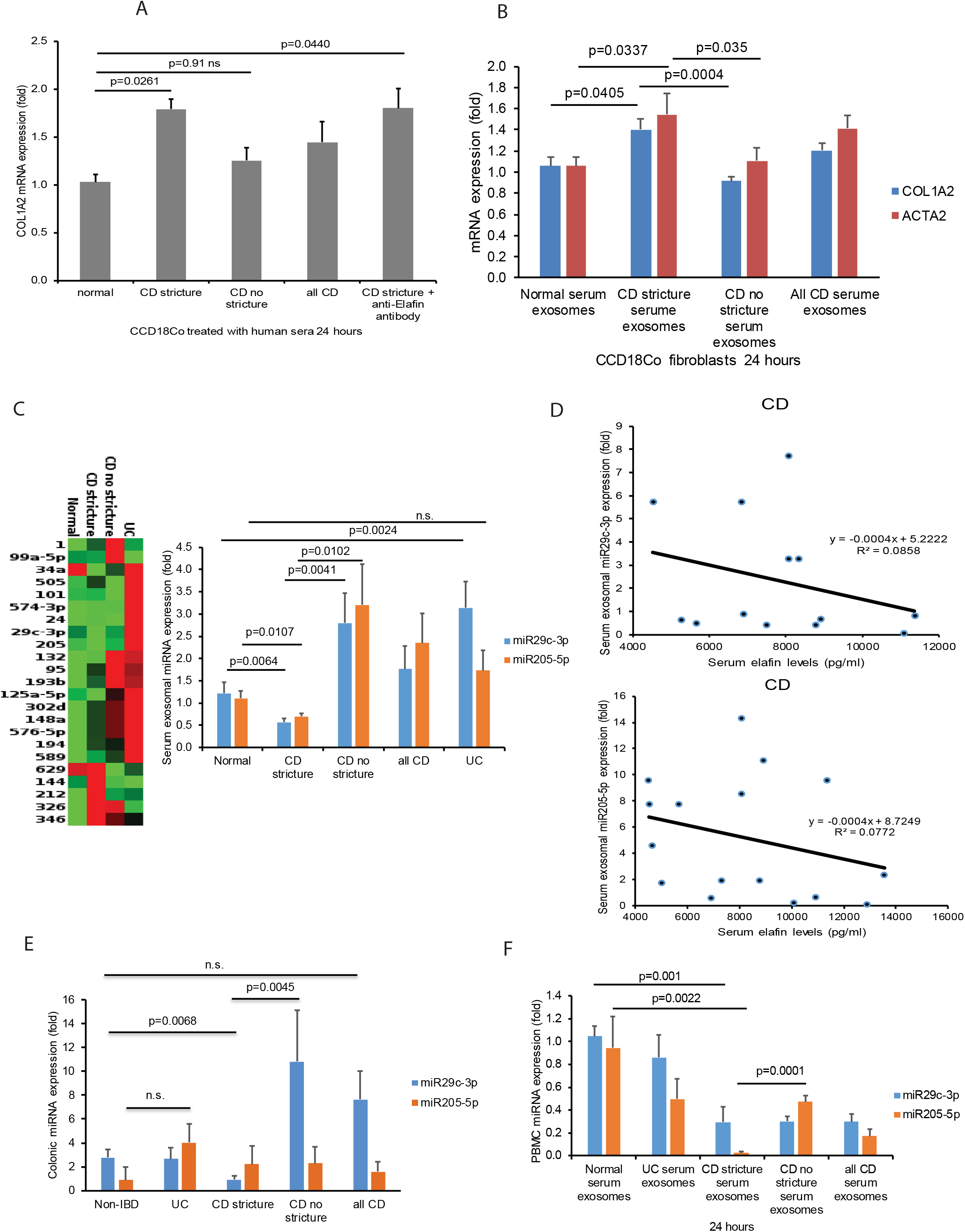
Circulating elafin levels are negatively correlated with serum exosomal anti-fibrogenic miRNAs. (A) The human intestinal fibroblasts were incubated with 100μl/ml of human sera in serum-free DMEM for 24 hours. (B) The human intestinal fibroblasts were incubated with 100μg/ml of human serum exosomes in serum-free DMEM for 24 hours. The collagen (COL1A2) mRNA expression was determined by real-time RT-PCR. Both *in vitro* experiments consisted of 12 patients per group. Multiple group comparison was done by one-way ANOVA. (C) A heatmap shows the miRNA expression in the serum exosomes. Red color indicates increased expression. Green color indicates decreased expression. Cutoff Ct is 35 cycles. Only detectable miRNAs are shown. The heatmap was generated online by Qiagen GeneGlobe Data Analysis Center. (D) Scatter plots show the negative correlation between serum elafin levels and serum exosomal miR29c-3p and miR205-5p expression in 24 CD patients. (E) Colonic miR29c-3p and miR205-5p expression in non-IBD, UC, stricturing CD, and non-stricturing CD patients. Colonic miR209c expression was reduced in stricturing CD patients. Multiple group comparison was done by one-way ANOVA. (F) PBMCs (10^5^ cells) from healthy normal patients were incubated with 100μg/ml of human serum exosomes from normal, UC, stricturing CD, and non-stricturing CD patients in serum-free DMEM for 24 hours. Intracellular miR29c-3p and miR205-5p expression were determined by real-time RT-PCR. Intracellular miR29c and miR205 expression in PBMCs was reduced after exposure to stricturing CD serum exosomes. Multiple group comparisons were done by one-way ANOVA. 12 patients per group.

### Serum elafin levels are negatively correlated with serum exosomal anti-fibrogenic miRNAs in CD patients

Since serum exosomes contain miRNAs, we utilized a PCR array and RT-PCR to profile the miRNA expression in serum exosomes in normal, UC, and CD patients with and without strictures. Out of all detectable miRNA, serum exosomal miR29c-3p and miR205-5p expression were reduced in stricturing CD patients (Figure 7C). Both of these two miRNAs are anti-fibrogenic (Gregory, Bert et al., 2008, Nijhuis, Biancheri et al., 2014). Interestingly, serum exosomal miR29c-3p and miR205-5p were inversely proportional to serum elafin levels (Figure 7D).

To find the sources of these miRNAs, we determined their expression in colonic and adipose tissues. Stricturing CD patients have significantly reduced colonic miR29c but not miR205 expression (Figure 7E). Mesenteric fat expresses miR205, but its expression was not dependent on CD/UC/stricture status or adipose elafin protein expression (Supplementary Figure 4B-C). miR29 was undetectable in all mesenteric fat tissues (data not shown).

We believe that circulating immune cells may express miRNAs. Although IBD patient-derived PBMCs were unavailable to us, we mimicked the environment of IBD by exposing PBMCs from normal patients to serum exosomes from normal, UC, and CD patients. Exposure to serum exosomes from stricturing CD patients caused significantly lower miR205 expression in PBMCs, compared to exposure to serum exosomes from non-stricturing CD patients and non-IBD patients (Figure 7F). Exposure to serum exosomes from either stricturing or non-stricturing CD patients led to similarly reduced miR29c expression in PBMCs (Figure 7F). These findings suggest that the reduced colonic miR29c expression and PBMC miR205 expression are associated with the low serum exosomal miR29c and miR205 in stricturing CD patients.

## Discussion

This report indicates elafin as a promising circulating protein biomarker for identifying intestinal strictures in CD patients. We understand that intestinal strictures are a complex phenomenon with many co-existing processes. We noticed that some of the non-stricturing CD patients also had high circulating elafin levels, leading to moderate accuracy when elafin alone was used in identifying stricturing CD patients. Elafin alone is not enough to indicate intestinal strictures accurately because the complexity of many clinical characteristics of the patients has not been considered (Isakov, Dotan et al., 2017, Waljee, Lipson et al., 2017). Traditional statistics are inadequate to recognize the specific pattern of intestinal stricture phenotypes. Machine learning, a branch of artificial intelligence, enhanced the accuracy of circulating elafin biomarker in identifying the presence of intestinal fibrosis among CD patients. We have used various combinations of available clinical data and elafin levels during the optimization step, so the current algorithm has the highest accuracy.

Two expert panels had attempted to establish consensus endpoints and criteria for diagnosis and response to therapy in stricturing CD (Danese, Bonovas et al., 2018, Rieder, Bettenworth et al., 2018). Most of the diagnostic approaches for intestinal strictures are based on radiological and endoscopic assessment, which are inherently inconvenient and expensive. Although serum elafin does not predict future development of intestinal strictures (data not shown), this minimally invasive circulating biomarker may be suitable for identifying high-risk stricturing CD patients for further evaluation.

With optimal cut-off points (8000pg/ml for CD and 18000pg/ml for UC), different elafin levels are associated with clinical disease activities with moderate accuracy for CD and UC patients (Supplementary Figure 1-2). However, the inclusion of machine learning could not improve their accuracy further (data not shown). It is unnecessary to develop an elafin test as a UC biomarker because CRP and FC are commonly used by clinicians, but both are not ideal for CD disease activity assessment (Costa, Mumolo et al., 2005, Menees, Powell et al., 2015, Mosli et al., 2015, Walsham & Sherwood, 2016, Yamaguchi, Takeuchi et al., 2016).

Our study shows small increases in serum elafin levels in IBD patients (Figure 1A). In contrast, Zhang *et al.* showed decreased elafin protein expression in the blood of Chinese IBD patients (Zhang et al., 2017, Zhang et al., 2016). We do not understand the discrepancy of these findings, but the positive correlation between increasing elafin levels and multiple cytokines, especially IFNγ and IL-5, may be associated with Th1 and Th2 immunity in CD and UC patients (Figure 1E and 3C-D). High circulating elafin levels reflect increased T cell-mediated immune activity. Similarly, circulating elafin levels are positively correlated with disease severity in patients with adaptive immune diseases (psoriasis and graft-vs-host disease) (Elgharib et al., 2019, Paczesny et al., 2010). Elafin-dependent IFNγ, IL-5, IL-12 p70, IL-2, IL-15, and TNFα are not associated with the presence of intestinal strictures, suggesting that fibrogenesis are not driven by these elafin-dependent cytokines (Figure 1E).

We did not fully understand the relationship between reduced colonic elafin expression and increased circulating elafin level in stricturing CD patients, but fibrotic intestine in stricturing CD patients may have impaired elafin expression because they are occupied by a significant number of fibroblasts. Fibroblasts are an insignificant producer of elafin (Protein Atlas). On the other hand, the increased elafin mRNA expression in mesenteric fat and its positive correlation with colonic fibrogenic gene mRNA expression suggests mesenteric fat as a significant source of circulating elafin in stricturing CD patients (Figure 6A-B). As elafin mRNA expression in mesenteric fat and colon showed a negative correlation (Figure 6C), we speculate that the increased mesenteric fat elafin production may be an attempt to compensate the down-regulated colonic elafin expression by raising circulating elafin protein level in the stricturing CD patients. Adipose tissue may be involved in stricture development (Buning et al., 2015, Erhayiem et al., 2011), but the functional significance of adipose elafin expression during the development of intestinal fibrosis still needs further investigation in the future.

As circulating elafin is not a direct driver of fibrogenesis (Figure 7A), the low serum exosomal miR29 and miR205 expression may be sufficient to explain the pro-fibrogenic effect of circulating exosomes from stricturing CD patients because they are well known for their anti-fibrogenic roles (Figure 7B) (Gregory et al., 2008, Nijhuis et al., 2014). Our evidence suggests that elafin-dependent serum exosomal miR29c expression is regulated by the intestine, while miR205 expression is regulated by circulating immune cells (Figure 7E-F).

High colonic elafin expression group had a small improvement in histology score (Supplementary Figure 3E-F), but this improvement was not sufficient to improve clinical disease activity in UC and CD patients (Supplementary Figure 3A-D). Likewise, circulating elafin does not affect intestinal injury in UC patients (Figure 3B). The increased endogenous elafin expression is insufficient to change the overall disease course among IBD patients. Although we are interested in discovering the functional roles of elafin in intestinal fibrosis, it is not feasible to evaluate the influence of endogenous elafin in the development of intestinal fibrosis in mice because they do not have the elafin gene.

A mouse study showed that elafin-overexpressing lactic acid bacteria are effective in inhibiting colitis (Motta, Bermudez-Humaran et al., 2012), suggesting that high intestinal levels of elafin are needed to ameliorate IBD. Our group is evaluating the potential of elafin as a therapeutic strategy against intestinal fibrosis because our preliminary experiments showed that elafin overexpression inhibited intestinal fibrosis in multiple mouse models (Trinitrobenzene sulfonic acid/TNBS, *Salmonella*, SAMP1/YitFc), while elafin peptide (in microgram per milliliter range) also reduced collagen (COL1A2) mRNA expression in human intestinal fibroblasts. Therefore, the increased circulating elafin level does not promote intestinal stricture development.

In summary, this study made a comprehensive measurement of elafin expression in circulation, colonic tissues, and mesenteric fat in non-IBD, UC, stricturing CD, and non-stricturing CD patients. Abnormal expression of elafin reflects the dynamics of local and systemic disease activities. Intestinal stricture is associated with increased circulating elafin levels, reduced intestinal elafin expression, and increased mesenteric fat elafin expression. Our study is the first to recognize elafin as a communication signal between mesenteric fat, blood, and intestine during stricture development. We utilized machine learning to integrate the elafin level and clinical data and developed an improved algorithm for indicating the presence of intestinal strictures in CD patients accurately.

## Contributors

JW, CO, LF, and YX performed the experiments. WH and AM contributed blood samples. DQS contributed colonic tissues and clinical data. HWK contributed to the conception and design, supervised the entire study, and wrote the manuscript. All authors revised the intellectual content and the manuscript critically. All authors have read and approved the final version of this manuscript. All agreed to be accountable for their respective aspects of contributions.

## Data Sharing Statement

The data and materials used in this study are available at Dr. Koon’s laboratory.

## Acknowledgment

We thank Prof. Charalabos Pothoulakis, MD for technical and financial assistance to this project. <u>The report is associated with a US provisional patent #62/650,981 filed on 3/31/2018</u>.

---

**Table 1**

**Baseline characteristics of serum samples.**

**Table 2**

**Disease locations and medications of IBD patients.**

**Table 3**

**Baseline characteristics of colonic and mesenteric fat samples.**

**Supplementary Figure 1**

**Circulating elafin is moderately accurate in indicating clinical disease activity in CD patients.**

(A) Serum elafin levels in CD patients with and without intestinal fistulas. (B-C) Prevalence, sensitivity, specificity, positive predictive value, negative predictive value, and odds ratio values of elafin test in indicating

(B) CD clinical remission and (C) moderate or severe CD clinical activity. (D) ROC curve with AUC value demonstrates the moderate accuracy of using elafin test for indicating CD clinical disease activity. Optimal cutoff point is 8000pg/ml.

**Supplementary Figure 2**

**Circulating elafin is moderately accurate in indicating clinical disease activity in UC patients.**

(A-B) Prevalence, sensitivity, specificity, positive predictive value, negative predictive value, and odds ratio values of elafin test in indicating (A) UC clinical remission and (B) moderate or severe UC clinical activity. (C) ROC curve with AUC value demonstrates the moderate accuracy of using elafin test for indicating UC clinical disease activity. Optimal cutoff point is 18000pg/ml.

**Supplementary Figure 3**

**Colonic elafin mRNA expression is negatively correlated with colonic injury in CD and UC patients.**

(A-B) Scatter plots show no significant correlation between clinical disease activity and colonic elafin mRNA expression in UC and CD patients. (C-D) Scatter plots show no significant correlation between clinical disease activity and colonic elafin protein expression in UC and CD patients. Simple Clinical Colitis Activity Score for UC patients. Harvey Bradshaw Index for CD patients. (E-F) Scatter plots show the weak negative correlation between colonic histology score and colonic elafin mRNA expression in UC and CD patients. The analysis included 26 UC patients and 29 CD patients.

**Supplementary Figure 4**

**Elafin mRNA expression is not correlated with miR205 expression in mesenteric fat in CD patients.**

(A) Immunohistochemistry of elafin in human mesenteric fat tissues at 400X magnification. Elafin (as shown by brown color) protein expression was strong around the adipocytes of mesenteric fat in stricturing CD patients. (B) Mesenteric fat miR205 expression. (C) Scatter plot shows no significant correlation between elafin mRNA expression and miR205 expression in the mesenteric fat in CD patients. The analysis included 19 CD patients.

## References

Bettenworth D, Nowacki TM, Cordes F, Buerke B, Lenze F (2016) Assessment of stricturing Crohn’s disease: Current clinical practice and future avenues. World J Gastroenterol 22: 1008–16

Buning C, von Kraft C, Hermsdorf M, Gentz E, Wirth EK, Valentini L, Haas V (2015) Visceral Adipose Tissue in Patients with Crohn’s Disease Correlates with Disease Activity, Inflammatory Markers, and Outcome. Inflamm Bowel Dis 21: 2590–7

Costa F, Mumolo MG, Ceccarelli L, Bellini M, Romano MR, Sterpi C, Ricchiuti A, Marchi S, Bottai M (2005) Calprotectin is a stronger predictive marker of relapse in ulcerative colitis than in Crohn’s disease. Gut 54: 364–8

D’Haens GR, Geboes K, Peeters M, Baert F, Penninckx F, Rutgeerts P (1998) Early lesions of recurrent Crohn’s disease caused by infusion of intestinal contents in excluded ileum. Gastroenterology 114: 262–7

Danese S, Bonovas S, Lopez A, Fiorino G, Sandborn WJ, Rubin DT, Kamm MA, Colombel JF, Sands BE, Vermeire S, Panes J, Rogler G, D’Haens G, Peyrin-Biroulet L (2018) Identification of Endpoints for Development of Antifibrosis Drugs for Treatment of Crohn’s Disease. Gastroenterology 155: 76–87

Elgharib I, Khashaba SA, Elsaid HH, Sharaf MM (2019) Serum elafin as a potential inflammatory marker in psoriasis. Int J Dermatol 58: 205–209

Erhayiem B, Dhingsa R, Hawkey CJ, Subramanian V (2011) Ratio of visceral to subcutaneous fat area is a biomarker of complicated Crohn’s disease. Clin Gastroenterol Hepatol 9: 684–687 e1

Flach CF, Eriksson A, Jennische E, Lange S, Gunnerek C, Lonnroth I (2006) Detection of elafin as a candidate biomarker for ulcerative colitis by whole-genome microarray screening. Inflamm Bowel Dis 12: 837–42

Gregory PA, Bert AG, Paterson EL, Barry SC, Tsykin A, Farshid G, Vadas MA, Khew-Goodall Y, Goodall GJ (2008) The miR-200 family and miR-205 regulate epithelial to mesenchymal transition by targeting ZEB1 and SIP1. Nat Cell Biol 10: 593–601

Hing TC, Ho S, Shih DQ, Ichikawa R, Cheng M, Chen J, Chen X, Law I, Najarian R, Kelly CP, Gallo RL, Targan SR, Pothoulakis C, Koon HW (2013) The antimicrobial peptide cathelicidin modulates Clostridium difficile-associated colitis and toxin A-mediated enteritis in mice. Gut 62: 1295–305

Ho S, Pothoulakis C, Koon HW (2013) Antimicrobial peptides and colitis. Curr Pharm Des 19: 40–7

Isakov O, Dotan I, Ben-Shachar S (2017) Machine Learning-Based Gene Prioritization Identifies Novel Candidate Risk Genes for Inflammatory Bowel Disease. Inflamm Bowel Dis 23: 1516–1523

Koon HW, Shih DQ, Chen J, Bakirtzi K, Hing TC, Law I, Ho S, Ichikawa R, Zhao D, Xu H, Gallo R, Dempsey P, Cheng G, Targan SR, Pothoulakis C (2011) Cathelicidin signaling via the Toll-like receptor protects against colitis in mice. Gastroenterology 141: 1852–63 e1-3

Kugathasan S, Denson LA, Walters TD, Kim MO, Marigorta UM, Schirmer M, Mondal K, Liu C, Griffiths A, Noe JD, Crandall WV, Snapper S, Rabizadeh S, Rosh JR, Shapiro JM, Guthery S, Mack DR, Kellermayer R, Kappelman MD, Steiner S et al. (2017) Prediction of complicated disease course for children newly diagnosed with Crohn’s disease: a multicentre inception cohort study. Lancet 389: 1710–1718

Lenti MV, Di Sabatino A (2019) Intestinal fibrosis. Mol Aspects Med 65: 100–109

Louis E, Collard A, Oger AF, Degroote E, Aboul Nasr El Yafi FA, Belaiche J (2001) Behaviour of Crohn’s disease according to the Vienna classification: changing pattern over the course of the disease. Gut 49: 777–82

Mao R, Kurada S, Gordon IO, Baker ME, Gandhi N, McDonald C, Coffey JC, Rieder F (2019) The Mesenteric Fat and Intestinal Muscle Interface: Creeping Fat Influencing Stricture Formation in Crohn’s Disease. Inflamm Bowel Dis 25: 421–426

Menees SB, Powell C, Kurlander J, Goel A, Chey WD (2015) A meta-analysis of the utility of C-reactive protein, erythrocyte sedimentation rate, fecal calprotectin, and fecal lactoferrin to exclude inflammatory bowel disease in adults with IBS. Am J Gastroenterol 110: 444–54

Mosli MH, Zou G, Garg SK, Feagan SG, MacDonald JK, Chande N, Sandborn WJ, Feagan BG (2015) C-Reactive Protein, Fecal Calprotectin, and Stool Lactoferrin for Detection of Endoscopic Activity in Symptomatic Inflammatory Bowel Disease Patients: A Systematic Review and Meta-Analysis. Am J Gastroenterol 110: 802–19; quiz 820

Motta JP, Bermudez-Humaran LG, Deraison C, Martin L, Rolland C, Rousset P, Boue J, Dietrich G, Chapman K, Kharrat P, Vinel JP, Alric L, Mas E, Sallenave JM, Langella P, Vergnolle N (2012) Food-grade bacteria expressing elafin protect against inflammation and restore colon homeostasis. Sci Transl Med 4: 158ra144

Nijhuis A, Biancheri P, Lewis A, Bishop CL, Giuffrida P, Chan C, Feakins R, Poulsom R, Di Sabatino A, Corazza GR, MacDonald TT, Lindsay JO, Silver AR (2014) In Crohn’s disease fibrosis-reduced expression of the miR-29 family enhances collagen expression in intestinal fibroblasts. Clin Sci (Lond) 127: 341–50

Olewicz-Gawlik A, Trzybulska D, Graniczna K, Kuznar-Kaminska B, Katulska K, Batura-Gabryel H, Frydrychowicz M, Danczak-Pazdrowska A, Mozer-Lisewska I (2017) Serum alarm antiproteases in systemic sclerosis patients. Hum Immunol 78: 559–564

Paczesny S, Braun TM, Levine JE, Hogan J, Crawford J, Coffing B, Olsen S, Choi SW, Wang H, Faca V, Pitteri S, Zhang Q, Chin A, Kitko C, Mineishi S, Yanik G, Peres E, Hanauer D, Wang Y, Reddy P et al. (2010) Elafin is a biomarker of graft-versus-host disease of the skin. Sci Transl Med 2: 13ra2

Rieder F, Bettenworth D, Ma C, Parker CE, Williamson LA, Nelson SA, van Assche G, Di Sabatino A, Bouhnik Y, Stidham RW, Dignass A, Rogler G, Taylor SA, Stoker J, Rimola J, Baker ME, Fletcher JG, Panes J, Sandborn WJ, Feagan BG et al. (2018) An expert consensus to standardise definitions, diagnosis and treatment targets for anti-fibrotic stricture therapies in Crohn’s disease. Aliment Pharmacol Ther 48: 347–357

Schmid M, Fellermann K, Fritz P, Wiedow O, Stange EF, Wehkamp J (2007) Attenuated induction of epithelial and leukocyte serine antiproteases elafin and secretory leukocyte protease inhibitor in Crohn’s disease. J Leukoc Biol 81: 907–15

Shaw L, Wiedow O (2011) Therapeutic potential of human elafin. Biochem Soc Trans 39: 1450–4

Sideri A, Bakirtzi K, Shih DQ, Koon HW, Fleshner P, Arsenescu R, Arsenescu V, Turner JR, Karagiannides I, Pothoulakis C (2015) Substance P mediates pro-inflammatory cytokine release form mesenteric adipocytes in Inflammatory Bowel Disease patients. Cell Mol Gastroenterol Hepatol 1: 420–432

Tai EK, Wu WK, Wong HP, Lam EK, Yu L, Cho CH (2007) A new role for cathelicidin in ulcerative colitis in mice. Exp Biol Med (Maywood) 232: 799–808

Tran DH, Wang J, Ha C, Ho W, Mattai SA, Oikonomopoulos A, Weiss G, Lacey P, Cheng M, Shieh C, Mussatto CC, Ho S, Hommes D, Koon HW (2017) Circulating cathelicidin levels correlate with mucosal disease activity in ulcerative colitis, risk of intestinal stricture in Crohn’s disease, and clinical prognosis in inflammatory bowel disease. BMC Gastroenterol 17: 63

Waljee AK, Lipson R, Wiitala WL, Zhang Y, Liu B, Zhu J, Wallace B, Govani SM, Stidham RW, Hayward R, Higgins PDR (2017) Predicting Hospitalization and Outpatient Corticosteroid Use in Inflammatory Bowel Disease Patients Using Machine Learning. Inflamm Bowel Dis 24: 45–53

Walsham NE, Sherwood RA (2016) Fecal calprotectin in inflammatory bowel disease. Clin Exp Gastroenterol 9: 21–9

Xu C, Ghali S, Wang J, Shih DQ, Ortiz C, Mussatto CC, Lee EC, Tran DH, Jacobs JP, Lagishetty V, Fleshner P, Robbins L, Vu M, Hing TC, McGovern DPB, Koon HW (2017) CSA13 inhibits colitis-associated intestinal fibrosis via a formyl peptide receptor like-1 mediated HMG-CoA reductase pathway. Sci Rep 7: 16351

Yamaguchi S, Takeuchi Y, Arai K, Fukuda K, Kuroki Y, Asonuma K, Takahashi H, Saruta M, Yoshida H (2016) Fecal calprotectin is a clinically relevant biomarker of mucosal healing in patients with quiescent ulcerative colitis. J Gastroenterol Hepatol 31: 93–8

Yoo JH, Ho S, Tran DH, Cheng M, Bakirtzi K, Kukota Y, Ichikawa R, Su B, Tran DH, Hing TC, Chang I, Shih DQ, Issacson RE, Gallo RL, Fiocchi C, Pothoulakis C, Koon HW (2015) Anti-fibrogenic effects of the anti-microbial peptide cathelicidin in murine colitis-associated fibrosis. Cell Mol Gastroenterol Hepatol 1: 55–74 e1

Zhang W, Teng G, Wu T, Tian Y, Wang H (2017) Expression and Clinical Significance of Elafin in Inflammatory Bowel Disease. Inflamm Bowel Dis 23: 2134–2141

Zhang W, Teng GG, Tian Y, Wang HH (2016) [Expression of elafin in peripheral blood in inflammatory bowel disease patients and its clinical significance]. Zhonghua Yi Xue Za Zhi 96: 1120–3

